# Protein Representation Learning via Knowledge Enhanced Primary Structure Modeling

**DOI:** 10.1101/2023.01.26.525795

**Authors:** Hong-Yu Zhou, Yunxiang Fu, Zhicheng Zhang, Cheng Bian, Yizhou Yu

## Abstract

Protein representation learning has primarily benefited from the remarkable development of language models (LMs). Accordingly, pre-trained protein models also suffer from a problem in LMs: a lack of factual knowledge. The recent solution models the relationships between protein and associated knowledge terms as the knowledge encoding objective. However, it fails to explore the relationships at a more granular level, i.e., the token level. To mitigate this, we propose Knowledge-exploited Auto-encoder for Protein (KeAP), which performs token-level knowledge graph exploration for protein representation learning. In practice, non-masked amino acids iteratively query the associated knowledge tokens to extract and integrate helpful information for restoring masked amino acids via attention. We show that KeAP can consistently outperform the previous counterpart on 9 representative downstream applications, sometimes surpassing it by large margins. These results suggest that KeAP provides an alternative yet effective way to perform knowledge enhanced protein representation learning. Code and models are available at https://github.com/RL4M/KeAP.

## 1 Introduction

The unprecedented success of AlphaFold (Jumper et al., 2021; Senior et al., 2020) has sparked the public’s interest in artificial intelligence-based protein science, which in turn promotes scientists to develop more powerful deep neural networks for protein. At present, a major challenge faced by researchers is how to learn generalized representation from a vast amount of protein data. An analogous problem also exists in natural language processing (NLP), while the recent development of big language models (Devlin et al., 2018; Brown et al., 2020) offers a viable solution: unsupervised pre-training with self-supervision. In practice, by viewing amino acids as language tokens, we can easily transfer existing unsupervised pre-training techniques from NLP to protein, and the effectiveness of these techniques has been verified in protein representation learning Rao et al. (2019); Alley et al. (2019); Elnaggar et al. (2021); Unsal et al. (2022).

However, as pointed out by (Peters et al., 2019; Zhang et al., 2019; Sun et al., 2020; Wang et al., 2021), pre-trained language models often suffer from a lack of factual knowledge. To alleviate similar problems appearing in protein models, Zhang et al. (2022) proposed OntoProtein that explicitly injects factual biological knowledge into the pre-trained model, leading to observable improvements on several downstream protein analysis tasks, such as amino acid contact prediction and proteinprotein interaction identification.

In practice, OntoProtein leverages the masked language modeling (MLM) (Devlin et al., 2018) and TransE (Bordes et al., 2013) objectives to perform structure and knowledge encoding, respectively. Specifically, the TransE objective is applied to triplets from knowledge graphs, where each triplet can be formalized as (*Protein, Relation, Attribute*). The relation and attribute terms described using natural language are from the gene ontologies (Ashburner et al., 2000) associated with protein. However, OntoProtein only models the relationships on top of the contextual representations of protein (averaging amino acid representations) and textual knowledge (averaging word representations), preventing it from exploring knowledge graphs at a more granular level, i.e., the token level.

We propose KeAP (**K**nowledge-**e**xploited **A**uto-encoder for **P**rotein) to perform knowledge enhanced protein representation learning. To address the granularity issue of OntoProtein, KeAP performs token-level protein-knowledge exploration using the cross-attention mechanism. Specifically, each amino acid iteratively queries each word from relation and attribute terms to extract useful, relevant information using QKV Attention (Vaswani et al., 2017). The extracted information is then integrated into the protein representation via residual learning (He et al., 2016). The training process is guided only by the MLM objective, while OntoProtein uses contrastive learning and masked modeling simultaneously. Moreover, we propose to explore the knowledge in a cascaded manner by first extracting information from relation terms and then from attribute terms, which performs more effective knowledge encoding.

KeAP has two advantages over OntoProtein (Zhang et al., 2022). First, KeAP explores knowledge graphs at a more granular level by applying cross-attention to sequences of amino acids and words from relation and attributes. Second, KeAP provides a neat solution for knowledge enhanced protein pre-training. The encoder-decoder architecture in KeAP can be trained using the MLM objective only (both contrastive loss and MLM are used in OntoProtein), making the whole framework easy to optimize and implement.

Experimental results verify the performance superiority of KeAP over OntoProtein. In Fig. 1, we fine-tune the pre-trained protein models on 9 downstream applications. We see that KeAP outperforms OntoProtein on all 9 tasks mostly by obvious margins, such as amino acid contact prediction and protein-protein interaction (PPI) identification. Compared to ProtBert, KeAP achieves better results on 8 tasks while performing comparably on protein stability prediction. In contrast, OntoProtein produces unsatisfactory results on homology, stability, and binding affinity prediction.

**Figure 1:**
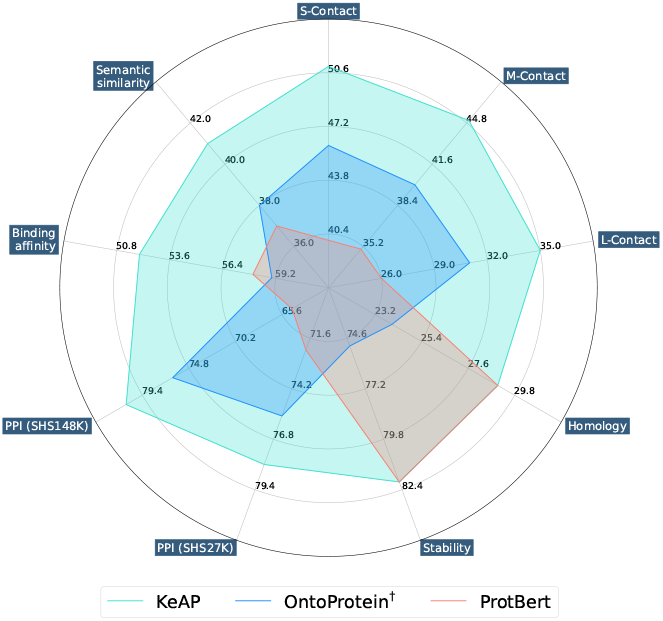
Transfer learning performance of ProtBert, OntoProtein, and our KeAP on downstream protein analysis tasks. S-, M-, and L-Contact stand for short-range, medium-range, and long-range contact prediction. PPI denotes the protein-protein interaction prediction. † means the model is trained with the full ProteinKG25.

## 2 Related Works

### 2.1 Representation Learning for Protein

How to learn generalized protein representation has recently become a hot topic in protein science, inspired by the widespread use of representation learning in language models (Devlin et al., 2018; Radford et al., 2019; Yang et al., 2019; Sarzynska-Wawer et al., 2021). Bepler & Berger (2018) introduced a multi-task protein representation learning framework, which obtains supervision signals from protein-protein structural similarity and individual amino acid contact maps. Due to a plethora of uncharacterized protein data, self-supervised pre-training (Alley et al., 2019; Rao et al., 2019) was proposed to directly learn representation from chains of amino acids, where tremendous and significant efforts were made to improve the pre-training result by scaling up the size of the model and dataset (Elnaggar et al., 2021; Rives et al., 2021; Vig et al., 2020; Rao et al., 2020; Yang et al., 2022; Nijkamp et al., 2022; Ferruz et al., 2022). In contrast, protein-related factual knowledge, providing abundant descriptive information for protein, has been long ignored and largely unexploited. OntoProtein (Zhang et al., 2022) first showed that we can improve the performance of pre-trained models on downstream tasks by explicitly injecting the factual biological knowledge associated with protein sequences into pre-training.

In practice, OntoProtein proposed to reconstruct masked amino acids while minimizing the embedding distance between contextual representations of protein and associated knowledge terms. One potential pitfall of this operation is that it fails to explore the relationships between protein and knowledge at a more granular level, i.e., the token level. In comparison, our KeAP overcomes this limitation by performing token-level protein-knowledge exploration via cross-attention modules.

### 2.2 Knowledge Enhanced Pre-trained Language Models

Knowledge integration has been treated as a reliable way to improve modern language models (Zhang et al., 2019; Sun et al., 2020; Liu et al., 2019; Vulic et al., 2020; Petroni et al., 2019; Roberts et al., 2020; Wang et al., 2021; Yao et al., 2019; Liu et al., 2020; He et al., 2020; Qin et al., 2021; Madani et al., 2020). Xie et al. (2016) proposed to perform end-to-end representation learning on triplets extracted from knowledge graphs, while Wang et al. (2021) further bridged the gap between knowledge graphs and pre-trained language models by treating entity descriptions as entity embeddings and jointly training the knowledge encoding (i.e., TransE (Bordes et al., 2013)) and MLM objectives.

Motivated by (Wang et al., 2021), OntoProtein (Zhang et al., 2022) used the MLM and TransE (Bordes et al., 2013) as two training objectives when learning protein representation on knowledge graphs. However, OntoProtein fails to explore the knowledge graphs at a more granular level, i.e., the token level. In contrast, KeAP performs token-level protein-knowledge exploration via the attention mechanism and provides a neat solution for knowledge enhanced protein pre-training by training an encoder-decoder architecture using only the MLM objective.

## 3 Methodologies

As shown in Fig. 2, for each triplet (*Protein, Relation, Attribute*) in the knowledge graphs, we apply random masking to the amino acid sequence while treating the relation and attribute terms as the associated factual knowledge. After feature extraction, representations of each triplet are sent to the protein decoder for reconstructing missing amino acids. The decoder model comprises *N* stacked protein-knowledge exploration (PiK) blocks. In each block, the amino acid representation iteratively queries, extracts, and integrates helpful, relevant information from word representations of associated relation and attribute terms in a cascaded manner. The resulting representation is used to restore the masked amino acids, guided by the MLM objective. After pre-training, the protein encoder can be transferred to various downstream tasks.

**Figure 2:**
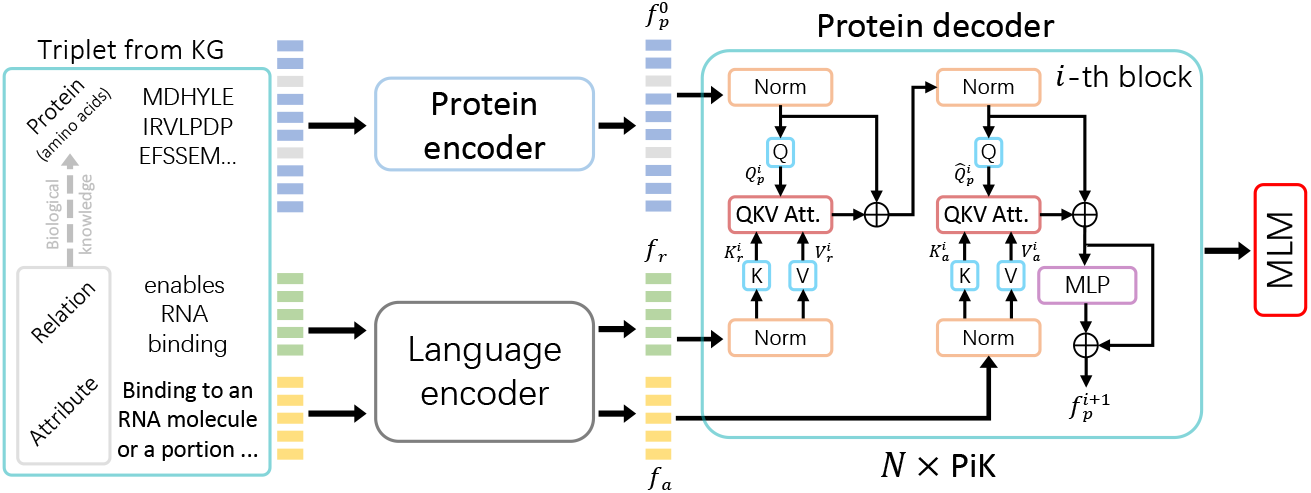
**Overview**. Given a triplet (*Protein, Relation, Attribute*) from the knowledge graph, KeAP randomly masks the input amino acid sequence and treats the relation and attribute terms as associated factual knowledge. Then, the triplet is passed to encoders and the output representations are regarded as the protein decoder’s inputs. The decoder asks each amino acid representation to iteratively query words from the associated knowledge terms in a cascaded manner, extracting and integrating helpful, relevant information for restoring masked amino acids. **PiK** and **MLM** stand for the protein-knowledge exploration block and masked language modeling, respectively.

### 3.1 Preliminary: Protein and Biological Knowledge

KeAP is trained on a knowledge graph that consists of about five million triplets from ProteinKG25 (Zhang et al., 2022). Each triplet is in the format of (*Protein, Relation, Attribute*). *Protein* can be viewed as a sequence of amino acids, while both *Relation* and *Attribute* are factual knowledge terms (denoted as gene ontologies in Zhang et al. (2022)) described using natural language. Specifically, *Relation* and *Attribute* provide knowledge in biology that is associated with *Protein*, such as molecular function, biological process, and cellular components.

During the training stage, each protein is passed to the protein encoder, resulting in the protein representation 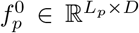. The superscript 0 is the layer index. *L_p_* denotes the length of the amino acid sequence. *D* stands for the feature dimension. In practice, the protein encoder has a BERT-like architecture (Devlin et al., 2018). Similarly, we forward the associated knowledge terms to the language encoder to obtain knowledge representations, i.e., *f_r_* ∈ ℝ^*L_r_*×*D*^ and *f_a_* ∈ ℝ^*L_a_*×*D*^. *L_r_* and *L_a_* denote the lengths of the relation and attribute terms, respectively. The reason for using two encoders is that we would like to extract domain-specific embeddings for protein and biological knowledge.

### 3.2 Token-level Protein-Knowledge Exploration

KeAP uses a surrogate task to perform knowledge enhanced pre-training, i.e., exploring knowledge graphs for protein primary structure modeling. In this way, KeAP asks the protein representation to be aware of what knowledge is helpful to masked protein modeling. In practice, we treat each amino acid as a query, while the words from associated relation and attribute terms are attended to as keys and values in order. Taking the *i*-th layer as an example, the inputs to the protein decoder include 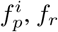, and *f_a_*. The relation representation *f_r_* is firstly queried by 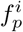 as the key and value:

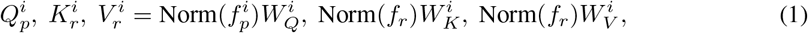

where 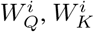, and 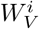 are learnable matrices. Norm stands for the layer normalization (Ba et al., 2016).

Then, QKV Attention (QKV-A) (Vaswani et al., 2017) is applied to 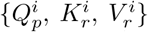, where representations of amino acids extract helpful, relevant knowledge from the latent word embeddings of the associated relation term. The obtained knowledge representation 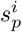 stores the helpful information for restoring missing amino acids. We then add up 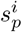 and 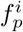 to integrate knowledge, resulting in the representation 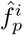.

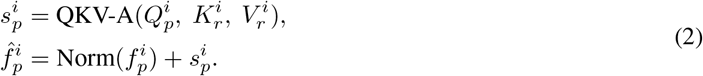

Next, we use 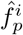 to query the attribute term. The whole query, extraction, and integration process is similar to that of the relation term:

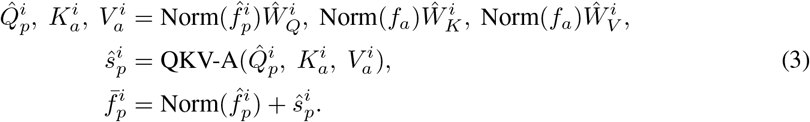

The resulting representation 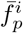 integrates the helpful, relevant biological knowledge that benefits the restoration of missing amino acids. We finally forward 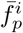 through a residual multi-layer perceptron (MLP) to obtain the output representation of the *i*-th PiK block, which also serves as the input to the *i*+1-th block.

### 3.3 Masked Language Modeling Objective

For each input protein, we randomly mask 20% amino acids in the sequence. Moreover, each masked amino acid has an 80% chance of being masked for prediction, a 10% chance of being replaced by a random amino acid, and a 10% chance of remaining unchanged. Suppose the number of masked amino acids is *M* and *x_j_* denotes the *j*-th amino acid. The training objective 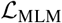 to be minimized is as follows:

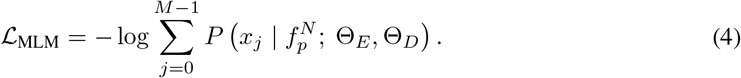

Θ_*E*_ and Θ_*D*_ denote the parameters of the protein encoder and decoder, respectively. We initialize the language encoder using PubMedBERT (Gu et al., 2021) and do not update it in the training phase.

## 4 Experiments and Analyses

In this section, we extensively evaluate the generalization ability of the learned protein representation by fine-tuning the pre-trained model on a wide range of downstream applications, including amino acid contact prediction, protein homology detection, protein stability prediction, proteinprotein interaction identification, protein-protein binding affinity prediction, and semantic similarity inference. Besides, we also provide ablations and failure analyses to facilitate the understanding of KeAP. Unless otherwise specified, we follow the pre-training and fine-tuning protocols used by OntoProtein (refer to the appendix for more details), such as training strategies and dataset split. The pre-trained models of ProtBert (Elnaggar et al., 2021), OntoProtein (Zhang et al., 2022), and our KeAP share the same number of network parameters. Average results are reported over three independent training runs.

### 4.1 Dataset for Pre-training

ProteinKG25 (Zhang et al., 2022) provides a knowledge graph that consists of approximately five million triplets, with nearly 600k protein, 50k attribute terms, and 31 relation terms included. The attribute and relation terms described using natural language are extracted from Gene Ontology^1^ which is the world’s largest source of information on the functions of genes and gene products (e.g., protein).

It is worth noting that we pre-train KeAP and report its experimental results in two different settings where the pre-training datasets differ. In the first setting, we remove 22,978 amino acid sequences that appear in downstream tasks. Accordingly, 396,993 triplets are removed from ProteinKG25, and the size of the new dataset is 91.87% of the original. In the second setting (denoted with ^†^), we use the same pre-training data as in OntoProtein (Zhang et al., 2022).

### 4.2 Amino Acid Contact Prediction

#### Overview

Given an input protein molecule that comprises a chain of amino acids, the protein model is asked to predict whether any two amino acids (from the same sequence) are in contact or not. To achieve this goal, our model outputs a probability contact matrix for each input protein, where the row and column numbers correspond to the indices of two amino acids. We performed experiments on the dataset collected and organized by (AlQuraishi, 2019; Rao et al., 2019). The evaluation metric is precision.

#### Baselines

Following (Zhang et al., 2022), we included six protein analysis models as baselines. Specifically, we used variants of LSTM (Hochreiter & Schmidhuber, 1997), ResNet (He et al., 2016), and Transformer (Vaswani et al., 2017) proposed by the TAPE benchmark (Rao et al., 2019). ProtBert (Elnaggar et al., 2021) is a 30-layer BERT-like model pre-trained on UniRef100 (Suzek et al., 2007; 2015). ESM-1b is a 33-layer transformer pre-trained on UR50/S, which uses the UniRef50 (Suzek et al., 2007; 2015) representative sequences. OntoProtein Zhang et al. (2022) is the most recent knowledge-based pre-training methodology.

#### Results

Table 1 presents the experimental results of amino acid contact prediction. In the first setting, we see that ProtBert and ESM-1b are two most competitive baselines. Specifically, KeAP outperforms ProtBert by large margins in short- (6 ≤ *seq* < 12), medium- (12 ≤ *seq* < 24), and long-range (seq ≥ 24) contact predictions. Compared to ESM-1b, KeAP is more advantageous in short-range prediction while achieving competitive results in medium- and long-range prediction tasks. In the second setting, we see that KeAP consistently surpasses OntoProtein regardless of the the distance between two amino acids. Particularly noteworthy is the fact that KeAP surpasses OntoProtein by large margins. Considering that OntoProtein also performs knowledge encoding, the clear performance advantages (6%) over OntoProtein demonstrate that KeAP provides a more competitive choice for knowledge enhanced protein representation learning. We believe the performance gains brought by KeAP can be attributed to the proposed token-level knowledge graph exploration methodology. This enables the pre-trained model to understand the knowledge context better, producing a better-contextualized protein representation for contact prediction than directly applying meaning pooling.

**Table 1:** Comparisons on amino acid contact prediction. **seq** indicates the distance (i.e., the number of amino acids) between two selected amino acids. **P@L**, **P@L/2**, **P@L/5** denote the precision scores calculated upon top L (i.e., L most likely contacts), top L/2, and top L/5 predictions, respectively. In each setting, the best results are bolded. Models with † are pre-trained using the full ProteinKG25.

| Methods | $6 \leq seq < 12$ | | | $12 \leq seq < 24$ | | | $24 \leq seq$ | | |
| --- | --- | --- | --- | --- | --- | --- | --- | --- | --- |
|  | P@L | P@L/2 | P@L/5 | P@L | P@L/2 | P@L/5 | P@L | P@L/2 | P@L/5 |
| LSTM | 0.26 | 0.36 | 0.49 | 0.20 | 0.26 | 0.34 | 0.20 | 0.23 | 0.27 |
| ResNet | 0.25 | 0.34 | 0.46 | 0.28 | 0.25 | 0.35 | 0.10 | 0.13 | 0.17 |
| Transformer | 0.28 | 0.35 | 0.46 | 0.19 | 0.25 | 0.33 | 0.17 | 0.20 | 0.24 |
| ProtBert | 0.30 | 0.40 | 0.52 | 0.27 | 0.35 | 0.47 | 0.20 | 0.26 | 0.34 |
| ESM-1b | 0.38 | 0.48 | 0.62 | 0.33 | 0.43 | **0.56** | 0.26 | 0.34 | **0.45** |
| KeAP | **0.41** | **0.51** | **0.63** | **0.36** | **0.45** | 0.54 | **0.28** | **0.35** | 0.43 |
| OntoProtein† | 0.37 | 0.46 | 0.57 | 0.32 | 0.40 | 0.50 | 0.24 | 0.31 | 0.39 |
| KeAP† | **0.41** | **0.52** | **0.62** | **0.36** | **0.46** | **0.57** | **0.29** | **0.37** | **0.46** |

### 4.3 Homology Detection and Stability Prediction

#### Overview of homology detection

Predicting the remote homology of protein can be formalized as a molecule-level classification task. Given a protein molecule as the input, we ask the homology detection model to predict the right protein fold type. In our case, there are 1,195 different types of protein folds, making it a quite challenging task. The data are from Hou et al. (2018) and we report average accuracy on the fold-level heldout set.

#### Overview of stability prediction

In this regression task, we aim to predict the intrinsic stability of the protein molecule, which measures the protein’s ability to maintain its fold under extreme conditions. In practice, high-accuracy stability prediction can benefit drug discovery, making it easier to construct stable protein molecules. We evaluate the model performance by calculating Spearman’s rank correlation scores on the whole test set Rocklin et al. (2017).

#### Baselines

Besides six approaches presented in Table 1, we add one more baseline Protein-Bert (Brandes et al., 2022), which pre-trains the protein model to restore missing amino acids and associated attribute annotations simultaneously.

#### Results

From Table 2, we see that the knowledge-based pre-training methodologies, i.e., Protein-Bert and OntoProtein, fail to display favorable results on both tasks. Zhang et al. (2022) claimed that the failure is due to the lack of sequence-level objectives in pre-training. in contrast, our KeAP performs on par with ProtBert on homology detection and protein stability prediction. We believe the success of KeAP can be partly attributed to the token-level knowledge graph exploration.

**Table 2:** Comparisons on protein homology detection and stability prediction. in each setting, the best results are bolded. Models with † are pre-trained using the full ProteinKG25.

| Methods | Homology | Stability |
| --- | --- | --- |
| LSTM | 0.26 | 0.69 |
| ResNet | 0.17 | 0.73 |
| Transformer | 0.21 | 0.73 |
| ProtBert | **0.29** | **0.82** |
| ProteinBert | 0.22 | 0.76 |
| ESM-1b | 0.11 | 0.77 |
| KeAP | **0.29** | **0.82** |
| OntoProtein<sup>†</sup> | 0.24 | 0.75 |
| KeAP<sup>†</sup> | **0.30** | **0.82** |

### 4.4 Protein-Protein Interaction Identification

#### Overview

Protein-protein interactions (PPI) refer to the physical contacts between two or more amino acid sequences. In this paper, we only study the two-protein cases, where a pair of protein molecules serve as the inputs. The goal is to predict the interaction type(s) of each protein pair. There are 7 types included in experiments, which are reaction, binding, post-translational modifications, activation, inhibition, catalysis, and expression. The problem of PPI prediction can be formalized as a multi-label classification problem. We perform experiments on SHS27K (Chen et al., 2019), SHS148K (Chen et al., 2019), and STRING (Lv et al., 2021). SHS27K and SHS148K can be regarded as two subsets of STRING, where protein with fewer than 50 amino acids or ≥ 40% sequence identity is excluded. Breadth-First Search (BFS) and Depth-First Search (DFS) are used to generate test sets from the aforementioned three datasets. F1 score is used as the default evaluation metric.

#### Baselines

Following Zhang et al. (2022), we introduce DPPI (Hashemifar et al., 2018), DNN-PPI (Li et al., 2018), PIPR (Chen et al., 2019), and GNN-PPI (Lv et al., 2021) as 4 more baselines in addition to ProtBert, ESM-1b, and OntoProtein.

#### Results

Experimental results are displayed in Table 3. In the first setting, we see that ESM-1b is the best performing baseline. Nonetheless, our KeAP still achieves the best average performance, outperforming ESM-1b on all three datasets by 1% at least. In the second setting, KeAP outperforms OntoProtein by about 4%, 3%, and 1% on SHS27K, SHS148K, and STRING, respectively. The trend of declining performance can be attributed to the increasing amount of fine-tuning data (from SHS27K to STRING) that reduces the impact of pre-training. Specifically, the advantages of KeAP are quite obvious on SHS27K which has the least number of protein, indicating the effectiveness of the protein representation from KeAP with limited fine-tuning data. As the amount of training data increases (from SHS27K to STRING), ProtBert and OntoProtein gradually display inferior performance, compared to GNN-PPI. In contrast, our KeAP still performs competitively and surpasses GNN-PPI by an obvious margin on BFS.

**Table 3:** Comparisons on PPI identification. Experiments were performed on three datasets, whose F1 scores are presented. The best results in each setting are bolded. Models with † are pre-trained using the full ProteinKG25.

| Methods | SHS27K |  |  | SHS148K |  |  | STRING |  |  |
| --- | --- | --- | --- | --- | --- | --- | --- | --- | --- |
|  | BFS | DFS | Avg | BFS | DFS | Avg | BFS | DFS | Avg |
| DNN-PPI | 48.09 | 54.34 | 51.22 | 57.40 | 58.42 | 57.91 | 53.05 | 64.94 | 59.00 |
| DPPI | 41.43 | 46.12 | 43.77 | 52.12 | 52.03 | 52.08 | 56.68 | 66.82 | 61.75 |
| PIPR | 44.48 | 57.80 | 51.14 | 61.83 | 63.98 | 62.91 | 55.65 | 67.45 | 61.55 |
| GNN-PPI | 63.81 | 74.72 | 69.27 | 71.37 | 82.67 | 77.02 | 78.37 | **91.07** | 84.72 |
| ProtBert | 70.94 | 73.36 | 72.15 | 70.32 | 78.86 | 74.59 | 67.61 | 87.44 | 77.53 |
| ESM-1b | 74.92 | **78.83** | 76.88 | **77.49** | 82.13 | 79.31 | 78.54 | 88.59 | 83.57 |
| KeAP | **78.58** | 77.54 | **78.06** | 77.22 | **84.74** | **80.98** | **81.44** | 89.77 | **85.61** |
| OntoProtein<sup>†</sup> | 72.26 | 78.89 | 75.58 | 75.23 | 77.52 | 76.38 | 76.71 | **91.45** | 84.08 |
| KeAP<sup>†</sup> | **79.17** | **79.77** | **79.47** | **75.67** | **83.04** | **79.36** | **80.78** | 89.02 | **84.90** |

### 4.5 Protein-Protein Binding Affinity Estimation

#### Overview

In this task, we aim to evaluate the ability of the protein representation to estimate the change of the binding affinity due to mutations of protein. In practice, each pair of protein is mapped to a real value (thus this is a regression task), indicating the binding affinity change. We follow (Unsal et al., 2022) to apply bayesian ridge regression to the result of the element-wise multiplication of representation extracted from pre-trained protein models for predicting the binding affinity. We used the SKEMPI dataset from (Moal & Fernández-Recio, 2012) and report the mean square error of 10-fold cross-validation.

#### Baselines

We include PIPR, ProtBert, ESM-1b, and OntoProtein as comparative baselines.

#### Results

Table 4 presents the experimental results on the binding affinity estimation task. We see that KeAP outperforms PIPR, ProtBert, and ESM-1b by substantial margins. Considering the region-level structural feature plays a vital role in this task (Unsal et al., 2022), we believe the obvious performance advantage of KeAP again verifies the effectiveness of the proposed token-level knowledge graph exploration methodology. Nonetheless, there is still a performance gap between our KeAP and ESM-1b, which may be attributed to the difference in network architecture.

**Table 4:** Comparisons on proteinprotein binding affinity prediction. The best result in each setting is bolded. ↓ means the lower the better. Models with † are pre-trained using the full ProteinKG25.

| Methods | Affinity (↓) |
| --- | --- |
| PIPR | 0.63 |
| ProtBert | 0.58 |
| ESM-1b | **0.50** |
| KeAP | 0.52 |
| OntoProtein<sup>†</sup> | 0.59 |
| KeAP<sup>†</sup> | 0.51 |

### 4.6 Semantic Similarity Inference

#### Overview

In this task, given two interacting protein molecules and their associated attribute terms, we first calculate the Manhattan Similarity^2^ between their representations. Then, we calculate the Lin Similarity between their associated attribute terms following instructions from Unsal et al. (2022). Finally, Spearman’s rank correlation is calculated between the Manhattan Similarity scores and Lin Similarity scores, where the Lin Similarity scores are treated as ground truths, and the Manhattan Similarity scores are regarded as predictions. Specifically, we divide protein attributes into three groups: molecular function (MF), biological process (BP), and cellular component (CC), and report the correlation scores for each group in Table 5.

**Table 5:** Comparisons on semantic similarity inference. The best results in each setting are bolded. Models with † are pre-trained using the full ProteinKG25.

| Methods | MF | BP | CC | Avg |
| --- | --- | --- | --- | --- |
| MSA Transformer | 0.38 | 0.31 | 0.30 | 0.33 |
| ProtT5-XL | **0.57** | 0.21 | **0.40** | 0.39 |
| ProtBert | 0.41 | 0.35 | 0.36 | 0.37 |
| ESM-1b | 0.38 | **0.42** | 0.37 | 0.39 |
| KeAP | 0.42 | 0.41 | 0.39 | **0.41** |
| OntoProtein<sup>†</sup> | **0.41** | 0.36 | 0.36 | 0.38 |
| KeAP<sup>†</sup> | **0.41** | **0.41** | **0.40** | **0.41** |

#### Baselines

In addition to ProtBert and OntoProtein, we introduce three powerful pretrained protein models for comparisons, which include MSA Transformer (Rao et al., 2021), ESM-1b (Rives et al., 2021), and ProtT5-XL (Elnaggar et al., 2021).

#### Results

Table 5 presents the similarity inference results. We see that KeAP achieves the best average result even compared to larger (with more parameters) protein models, such as ESM-1b and ProtT5-XL. Specifically, ProtT5-XL produces the best performance on MF and CC, while ESM-1b performs the best on BP. Compared to ESM-1b and ProtT5-XL, our KeAP gets 2nd place on MF and achieves the highest score on CC. These results demonstrate the potential of KeAP in outperforming big protein models, and it would be interesting if KeAP could be integrated into bigger models. Again, KeAP outperforms OntoProtein by substantial margins.

## 5 Ablation and Discussion

We present ablation study results in Tables 6, 7, and 8, where we investigate the impacts of using different pre-training datasets, different mask ratios, and different knowledge integration strategies, respectively. Table 9 presents the performance on three tasks from Rao et al. (2019). SS-Q3 and SS-Q8 are two secondary structure prediction tasks (Klausen et al., 2019; Cuff & Barton, 1999) with different numbers of local structures. Fluorescence is a regression task, where the model is asked to predict the log-fluorescence intensity of each protein.

**Table 6:** Performance changes when removing the proteins that appear in downstream tasks from the pre-training dataset (ProteinKG25). Average results are reported on contact prediction and PPI tasks.

|  | Contact | Homology | Stability | PPI | Affinity | Semantic Similarity |
| --- | --- | --- | --- | --- | --- | --- |
| Performance changes | -1% | -1% | 0% | +1% | -1% | 0% |

**Table 7:** Ablations of mask ratios. Medium-range P@L/2 results are reported for contact prediction.

| Ratios | Contact | Homology | PPI |
| --- | --- | --- | --- |
| 15% | 0.44 | 0.27 | 76.72 |
| 20% | 0.45 | 0.29 | 78.06 |
| 25% | 0.46 | 0.28 | 77.34 |

**Table 8:** Investigation of knowledge exploitation strategies.

| Strategies | Contact |
| --- | --- |
| KeAP | 0.45 |
| - Cascaded | 0.44 |
| - PiK | 0.38 |
| + Tri. Match | 0.45 |

**Table 9:** Failure case analysis. We report the results on three tasks from (Rao et al., 2019).

| Methods | SS-Q3 | SS-Q8 | Fluorescence |
| --- | --- | --- | --- |
| ProtBert | 0.81 | 0.67 | 0.67 |
| ESM-1b | - | 0.71 | 0.65 |
| KeAP | 0.82 | 0.67 | 0.65 |
| OntoProtein<sup>†</sup> | 0.82 | 0.68 | 0.67 |
| KeAP<sup>†</sup> | 0.82 | 0.68 | 0.67 |

### 5.1 Ablation Study

Table 6 presents the performance drops when we remove the proteins that appear in downstream tasks from ProteinKG25. We see that the pre-trained model shows slightly worse results on three out of six downstream tasks: contact prediction, homology detection, and affinity prediction, while performing competitively on PPI, stability prediction, and semantic similarity inference. These comparisons imply that we may not have to worry too much about the consequences of data leakage. We will continue to investigate this problem in future work.

As shown in Table 7, the 20% mask ratio performs the best on two (homology and PPI) of the three downstream tasks, which is the primary reason that we choose 20% as the default mask ratio. It is interesting that a larger ratio (i.e., 25%) leads to better performance on the contact prediction task. We leave the exploration of larger mask ratios for future work.

In addition to the mask ratio, we also study the impacts of using different knowledge exploitation strategies. First of all, removing the cascaded exploitation strategy (presented as - Cascaded in Table 8) results in a 1-percent performance drop on contact prediction, implying exploring the factual knowledge in a cascaded manner is a more effective choice. Then, we remove the proposed protein-knowledge exploration block (denoted as - PiK in Table 8), which means KeAP is simplified to an auto-encoder trained with the MLM objective. We find that this activity leads to an 7-percent performance drop on the contact prediction task, reflecting the necessity of incorporating knowledge into protein pre-training. Besides, we add a Triplet Matching training objective (appeared as - Tri. Match in Table 8) to KeAP, where we randomly replace the associated attribute term with a different one and train the model to tell whether the input triplet is matched or not. The idea is similar to that of the knowledge-aware contrastive learning proposed by OntoProtein (Zhang et al., 2022), which is learning the knowledge-aware protein representation. From Table 8, we see that adding the matching objective does not bring performance improvements to KeAP, indicating that our proposed exploration strategy for knowledge graphs may already master the information introduced by the Triplet Matching objective.

### 5.2 Failure Case Analysis

We report the experimental results on three tasks from Rao et al. (2019), where KeAP performs on par with or worse than ProtBert, ESM-1b, or OntoProtein. Specifically, in SS-Q3 and SS-Q8, the model is asked to predict the secondary structure of each amino acid, which heavily relies on the local information contained in the protein representation. We think the non-significant performance of KeAP is due to the lack of the incorporation of local details when performing the knowledge encoding. Similarly, on the Fluorescence task, KeAP also fails to achieve observable progress when asked to distinguish very similar protein molecules. Considering the same issues also exist in ProtBert and OntoProtein, we believe it is necessary to pay more attention to how to improve the performance on local prediction tasks by integrating more local information during the pre-training stage. We will continue to explore this issue in the future.

## 6 Conclusion and Future Work

We present KeAP, a new knowledge enhanced representation learning methodology for protein. In practice, KeAP performs token-level knowledge graph exploration using cross-modal attention. KeAP provides a neat solution for knowledge enhanced protein pre-training. The encoder-decoder architecture in KeAP can be trained using the MLM objective only (both contrastive loss and MLM are used in OntoProtein), making the whole framework easy to optimize and implement. KeAP outperforms the previous knowledge enhanced counterpart on 9 downstream applications, sometimes by substantial margins, demonstrating the performance superiority of KeAP. In the future, we will investigate how to deploy KeAP on specific applications where the factual knowledge makes a greater impact.

## A Appendix

### A.1 Hyper-parameters for Fine-tuning

The hyper-parameters for fine-tuning are provided in Table 10. Specifically, we follow the hyperparameter settings in GNN-PPI (Lv et al., 2021) for PPI prediction. For protein binding affinity prediction and semantic similarity inference, we follow the fine-tuning configurations in PROBE (Unsal et al., 2022).

**Table 10:** Hyper-parameters for fine-tuning.

| Task | Epoch | Batch size | Warmup ratio | Learning rate | Freeze Bert | Optimizer |
| --- | --- | --- | --- | --- | --- | --- |
| Contact | 5 | 8 | 0.08 | 3e-5 | false | AdamW |
| Homology | 10 | 32 | 0.08 | 4e-5 | false | AdamW |
| Stability | 5 | 64 | 0.08 | 1e-5 | false | AdamW |
| SS-Q3 | 5 | 32 | 0.08 | 3e-5 | false | AdamW |
| SS-Q8 | 5 | 32 | 0.08 | 3e-5 | false | AdamW |
| Fluorescence | 15 | 64 | 0.10 | 1e-3 | true | Adam |

### A.2 Probability Contact Maps

In Fig. 3, we present two randomly picked samples for visual analysis, where KeAP is able to detect contacts missed by ProtBert and OntoProtein with high confidence.

**Figure 3:**
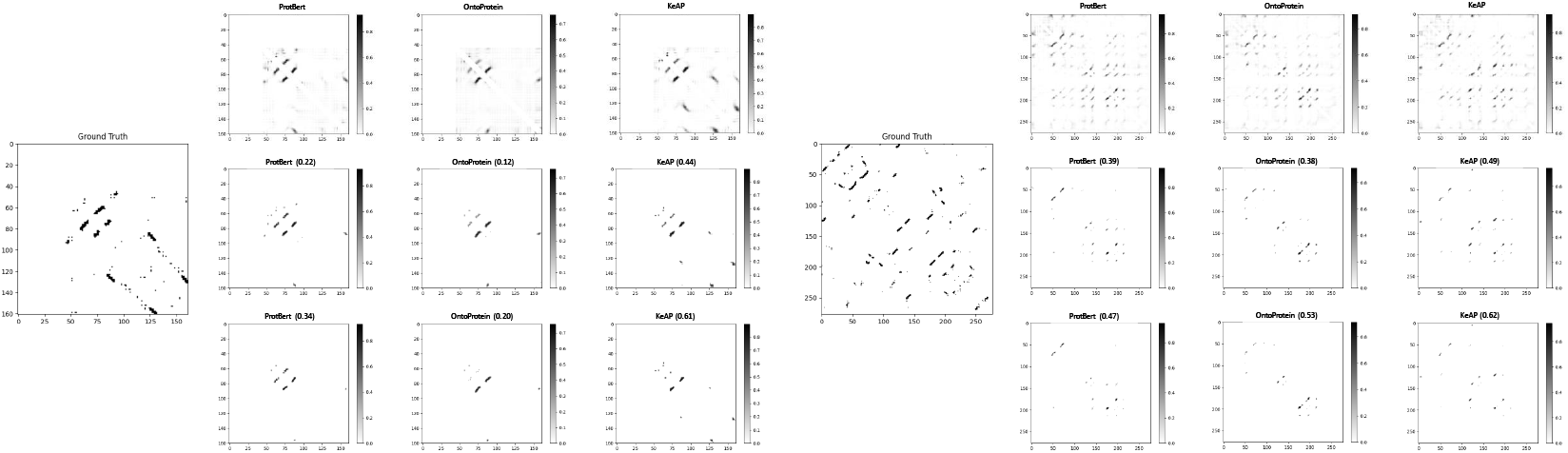
Ground truths and predicted probability contact maps. We compare the predictions of ProtBert and OntoProtein with ours, where the 1st, 2nd, and 3rd rows include the all, top L, and top L/2 predictions, respectively. For top L and L/2 predictions, the precision scores are reported. More examples are in the appendix.

## Footnotes

1 http://geneontology.org/

2 Manhattan Similarity = 1 - Manhattan Distance (normalized).

## References

Ethan C Alley, Grigory Khimulya, Surojit Biswas, Mohammed AlQuraishi, and George M Church. Unified rational protein engineering with sequence-based deep representation learning. Nature methods, 16(12):1315–1322, 2019.

Mohammed AlQuraishi. ProteinNet: a standardized data set for machine learning of protein structure. BMC bioinformatics, 20(1):1–10, 2019.

Michael Ashburner, Catherine A Ball, Judith A Blake, David Botstein, Heather Butler, J Michael Cherry, Allan P Davis, Kara Dolinski, Selina S Dwight, Janan T Eppig, et al. Gene Ontology: tool for the unification of biology. Nature genetics, 25(1):25–29, 2000.

Jimmy Lei Ba, Jamie Ryan Kiros, and Geoffrey E Hinton. Layer normalization. arXiv preprint arXiv:1607.06450, 2016.

Tristan Bepler and Bonnie Berger. Learning protein sequence embeddings using information from structure. In International Conference on Learning Representations, 2018.

Antoine Bordes, Nicolas Usunier, Alberto Garcia-Duran, Jason Weston, and Oksana Yakhnenko. Translating embeddings for modeling multi-relational data. Advances in neural information processing systems, 26, 2013.

Nadav Brandes, Dan Ofer, Yam Peleg, Nadav Rappoport, and Michal Linial. ProteinBERT: A universal deep-learning model of protein sequence and function. Bioinformatics, 38(8):2102–2110, 2022.

Tom Brown, Benjamin Mann, Nick Ryder, Melanie Subbiah, Jared D Kaplan, Prafulla Dhariwal, Arvind Neelakantan, Pranav Shyam, Girish Sastry, Amanda Askell, et al. Language models are few-shot learners. Advances in neural information processing systems, 33:1877–1901, 2020.

Muhao Chen, Chelsea J-T Ju, Guangyu Zhou, Xuelu Chen, Tianran Zhang, Kai-Wei Chang, Carlo Zaniolo, and Wei Wang. Multifaceted protein–protein interaction prediction based on siamese residual RCNN. Bioinformatics, 35(14):i305–i314, 2019.

James A Cuff and Geoffrey J Barton. Evaluation and improvement of multiple sequence methods for protein secondary structure prediction. Proteins: Structure, Function, and Bioinformatics, 34 (4):508–519, 1999.

Jacob Devlin, Ming-Wei Chang, Kenton Lee, and Kristina Toutanova. BERT: Pre-training of deep bidirectional transformers for language understanding. arXiv preprint arXiv:1810.04805, 2018.

Ahmed Elnaggar, Michael Heinzinger, Christian Dallago, Ghalia Rehawi, Yu Wang, Llion Jones, Tom Gibbs, Tamas Feher, Christoph Angerer, Martin Steinegger, et al. ProtTrans: Towards cracking the language of lifes code through self-supervised deep learning and high performance computing. IEEE Transactions on Pattern Analysis and Machine Intelligence, 2021.

Noelia Ferruz, Steffen Schmidt, and Birte Höcker. ProtGPT2 is a deep unsupervised language model for protein design. Nature communications, 13(1):1–10, 2022.

Yu Gu, Robert Tinn, Hao Cheng, Michael Lucas, Naoto Usuyama, Xiaodong Liu, Tristan Naumann, Jianfeng Gao, and Hoifung Poon. Domain-specific language model pretraining for biomedical natural language processing. ACM Transactions on Computing for Healthcare (HEALTH), 3(1): 1–23, 2021.

Somaye Hashemifar, Behnam Neyshabur, Aly A Khan, and Jinbo Xu. Predicting protein–protein interactions through sequence-based deep learning. Bioinformatics, 34(17):i802–i810, 2018.

Bin He, Di Zhou, Jinghui Xiao, Xin Jiang, Qun Liu, Nicholas Jing Yuan, and Tong Xu. BERT-MK: Integrating graph contextualized knowledge into pre-trained language models. In Findings of the Association for Computational Linguistics: EMNLP 2020, pp. 2281–2290, 2020.

Kaiming He, Xiangyu Zhang, Shaoqing Ren, and Jian Sun. Deep residual learning for image recognition. In Proceedings of the IEEE conference on computer vision and pattern recognition, pp. 770–778, 2016.

Sepp Hochreiter and Jürgen Schmidhuber. Long short-term memory. Neural computation, 9(8): 1735–1780, 1997.

Jie Hou, Badri Adhikari, and Jianlin Cheng. DeepSF: deep convolutional neural network for mapping protein sequences to folds. Bioinformatics, 34(8):1295–1303, 2018.

John M. Jumper, Richard Evans, Alexander Pritzel, Tim Green, Michael Figurnov, Olaf Ronneberger, Kathryn Tunyasuvunakool, Russ Bates, Augustin Zídek, Anna Potapenko, Alex Bridgland, Clemens Meyer, Simon A A Kohl, Andy Ballard, Andrew Cowie, Bernardino Romera-Paredes, Stanislav Nikolov, Rishub Jain, Jonas Adler, Trevor Back, Stig Petersen, David A. Reiman, Ellen Clancy, Michal Zielinski, Martin Steinegger, Michalina Pacholska, Tamas Berghammer, Sebastian Bodenstein, David Silver, Oriol Vinyals, Andrew W. Senior, Koray Kavukcuoglu, Pushmeet Kohli, and Demis Hassabis. Highly accurate protein structure prediction with alphafold. Nature, 596:583 – 589, 2021.

Michael Schantz Klausen, Martin Closter Jespersen, Henrik Nielsen, Kamilla Kjaergaard Jensen, Vanessa Isabell Jurtz, Casper Kaae Soenderby, Morten Otto Alexander Sommer, Ole Winther, Morten Nielsen, Bent Petersen, et al. NetSurfP-2.0: Improved prediction of protein structural features by integrated deep learning. Proteins: Structure, Function, and Bioinformatics, 2019.

Hang Li, Xiu-Jun Gong, Hua Yu, and Chang Zhou. Deep neural network based predictions of protein interactions using primary sequences. Molecules, 23(8):1923, 2018.

Nelson F Liu, Matt Gardner, Yonatan Belinkov, Matthew E Peters, and Noah A Smith. Linguistic knowledge and transferability of contextual representations. In Proceedings of the 2019 Conference of the North American Chapter of the Association for Computational Linguistics: Human Language Technologies, Volume 1 (Long and Short Papers), pp. 1073–1094, 2019.

Weijie Liu, Peng Zhou, Zhe Zhao, Zhiruo Wang, Qi Ju, Haotang Deng, and Ping Wang. K-bert: Enabling language representation with knowledge graph. In Proceedings of the AAAI Conference on Artificial Intelligence, volume 34, pp. 2901–2908, 2020.

Guofeng Lv, Zhiqiang Hu, Yanguang Bi, and Shaoting Zhang. Learning Unknown from Correlations: Graph neural network for inter-novel-protein interaction prediction. In Proceedings of the Thirtieth International Joint Conference on Artificial Intelligence, pp. 3677–3683, 2021.

Ali Madani, Bryan McCann, Nikhil Naik, Nitish Shirish Keskar, Namrata Anand, Raphael R Eguchi, Po-Ssu Huang, and Richard Socher. Progen: Language modeling for protein generation. arXiv preprint arXiv:2004.03497, 2020.

Iain H Moal and Juan Fernández-Recio. SKEMPI: a structural kinetic and energetic database of mutant protein interactions and its use in empirical models. Bioinformatics, 28(20):2600–2607, 2012.

Erik Nijkamp, Jeffrey Ruffolo, Eli N Weinstein, Nikhil Naik, and Ali Madani. ProGen2: exploring the boundaries of protein language models. arXiv preprint arXiv:2206.13517, 2022.

Matthew E. Peters, Mark Neumann, IV RobertL. Logan, Roy Schwartz, Vidur Joshi, Sameer Singh, and Noah A. Smith. Knowledge enhanced contextual word representations. In EMNLP, 2019.

Fabio Petroni, Tim Rocktäschel, Sebastian Riedel, Patrick Lewis, Anton Bakhtin, Yuxiang Wu, and Alexander Miller. Language models as knowledge bases? In Proceedings of the 2019 Conference on Empirical Methods in Natural Language Processing and the 9th International Joint Conference on Natural Language Processing (EMNLP-IJCNLP), pp. 2463–2473, 2019.

Yujia Qin, Yankai Lin, Ryuichi Takanobu, Zhiyuan Liu, Peng Li, Heng Ji, Minlie Huang, Maosong Sun, and Jie Zhou. ERICA: Improving entity and relation understanding for pre-trained language models via contrastive learning. In Joint Conference of the 59th Annual Meeting of the Association for Computational Linguistics and the 11th International Joint Conference on Natural Language Processing, ACL-IJCNLP 2021, pp. 3350–3363. Association for Computational Linguistics (ACL), 2021.

Alec Radford, Jeffrey Wu, Rewon Child, David Luan, Dario Amodei, Ilya Sutskever, et al. Language models are unsupervised multitask learners. OpenAI blog, 1(8):9, 2019.

Roshan Rao, Nicholas Bhattacharya, Neil Thomas, Yan Duan, Peter Chen, John Canny, Pieter Abbeel, and Yun Song. Evaluating protein transfer learning with TAPE. Advances in neural information processing systems, 32, 2019.

Roshan Rao, Joshua Meier, Tom Sercu, Sergey Ovchinnikov, and Alexander Rives. Transformer protein language models are unsupervised structure learners. In International Conference on Learning Representations, 2020.

Roshan M Rao, Jason Liu, Robert Verkuil, Joshua Meier, John Canny, Pieter Abbeel, Tom Sercu, and Alexander Rives. MSA transformer. In International Conference on Machine Learning, pp. 8844–8856. PMLR, 2021.

Alexander Rives, Joshua Meier, Tom Sercu, Siddharth Goyal, Zeming Lin, Jason Liu, Demi Guo, Myle Ott, C Lawrence Zitnick, Jerry Ma, et al. Biological structure and function emerge from scaling unsupervised learning to 250 million protein sequences. Proceedings of the National Academy of Sciences, 118(15):e2016239118, 2021.

Adam Roberts, Colin Raffel, and Noam Shazeer. How much knowledge can you pack into the parameters of a language model? In Proceedings of the 2020 Conference on Empirical Methods in Natural Language Processing (EMNLP), pp. 5418–5426, 2020.

Gabriel J Rocklin, Tamuka M Chidyausiku, Inna Goreshnik, Alex Ford, Scott Houliston, Alexander Lemak, Lauren Carter, Rashmi Ravichandran, Vikram K Mulligan, Aaron Chevalier, et al. Global analysis of protein folding using massively parallel design, synthesis, and testing. Science, 357 (6347):168–175, 2017.

Justyna Sarzynska-Wawer, Aleksander Wawer, Aleksandra Pawlak, Julia Szymanowska, Izabela Stefaniak, Michal Jarkiewicz, and Lukasz Okruszek. Detecting formal thought disorder by deep contextualized word representations. Psychiatry Research, 304:114135, 2021.

Andrew W. Senior, Richard Evans, John M. Jumper, James Kirkpatrick, L. Sifre, Tim Green, Chongli Qin, Augustin Zídek, Alexander W. R. Nelson, Alex Bridgland, Hugo Penedones, Stig Petersen, Karen Simonyan, Steve Crossan, Pushmeet Kohli, David T. Jones, David Silver, Koray Kavukcuoglu, and Demis Hassabis. Improved protein structure prediction using potentials from deep learning. Nature, 577:706–710, 2020.

Yu Sun, Shuohuan Wang, Yukun Li, Shikun Feng, Hao Tian, Hua Wu, and Haifeng Wang. ERNIE 2.0: A continual pre-training framework for language understanding. In Proceedings of the AAAI Conference on Artificial Intelligence, volume 34, pp. 8968–8975, 2020.

Baris E Suzek, Hongzhan Huang, Peter McGarvey, Raja Mazumder, and Cathy H Wu. Uniref: comprehensive and non-redundant uniprot reference clusters. Bioinformatics, 23(10):1282–1288, 2007.

Baris E Suzek, Yuqi Wang, Hongzhan Huang, Peter B McGarvey, Cathy H Wu, and UniProt Consortium. Uniref clusters: a comprehensive and scalable alternative for improving sequence similarity searches. Bioinformatics, 31(6):926–932, 2015.

Serbulent Unsal, Heval Atas, Muammer Albayrak, Kemal Turhan, Aybar C Acar, and Tunca Doğan. Learning functional properties of proteins with language models. Nature Machine Intelligence, 4 (3):227–245, 2022.

Ashish Vaswani, Noam Shazeer, Niki Parmar, Jakob Uszkoreit, Llion Jones, Aidan N Gomez, Łukasz Kaiser, and Illia Polosukhin. Attention is all you need. Advances in neural information processing systems, 30, 2017.

Jesse Vig, Ali Madani, Lav R Varshney, Caiming Xiong, Nazneen Rajani, et al. BERTology Meets Biology: Interpreting attention in protein language models. In International Conference on Learning Representations, 2020.

Ivan Vulić, Edoardo Maria Ponti, Robert Litschko, Goran Glavaš, and Anna Korhonen. Probing pretrained language models for lexical semantics. In Proceedings of the 2020 Conference on Empirical Methods in Natural Language Processing (EMNLP), pp. 7222–7240, 2020.

Xiaozhi Wang, Tianyu Gao, Zhaocheng Zhu, Zhengyan Zhang, Zhiyuan Liu, Juanzi Li, and Jian Tang. KEPLER: A unified model for knowledge embedding and pre-trained language representation. Transactions of the Association for Computational Linguistics, 9:176–194, 2021.

Ruobing Xie, Zhiyuan Liu, Jia Jia, Huanbo Luan, and Maosong Sun. Representation learning of knowledge graphs with entity descriptions. In Proceedings of the AAAI Conference on Artificial Intelligence, volume 30, 2016.

Kevin K Yang, Alex Xijie Lu, and Nicolo Fusi. Convolutions are competitive with transformers for protein sequence pretraining. In ICLR2022 Machine Learning for Drug Discovery, 2022.

Zhilin Yang, Zihang Dai, Yiming Yang, Jaime Carbonell, Russ R Salakhutdinov, and Quoc V Le. XLNet: Generalized autoregressive pretraining for language understanding. Advances in neural information processing systems, 32, 2019.

Liang Yao, Chengsheng Mao, and Yuan Luo. KG-BERT: Bert for knowledge graph completion. arXiv preprint arXiv:1909.03193, 2019.

Ningyu Zhang, Zhen Bi, Xiaozhuan Liang, Siyuan Cheng, Haosen Hong, Shumin Deng, Jiazhang Lian, Qiang Zhang, and Huajun Chen. OntoProtein: Protein pretraining with gene ontology embedding. International Conference on Learning Representations, 2022.

Zhengyan Zhang, Xu Han, Zhiyuan Liu, Xin Jiang, Maosong Sun, and Qun Liu. ERNIE: Enhanced language representation with informative entities. In Proceedings of the 57th Annual Meeting of the Association for Computational Linguistics, pp. 1441–1451, 2019.

